# Genomic aberration based molecular signatures efficiently characterize homologous recombination deficiency in prostate cancer

**DOI:** 10.1101/681114

**Authors:** Zsofia Sztupinszki, Miklos Diossy, Marcin Krzystanek, Judit Borcsok, Mark Pomerantz, Viktoria Tisza, Sandor Spisak, Orsolya Rusz, István Csabai, Matthew Freedman, Zoltan Szallasi

## Abstract

**Background:** Prostate cancers with mutations in genes involved in homologous recombination (HR), most commonly BRCA2, respond favorably to PARP inhibition and platinum-based chemotherapy. It is not clear, however, whether other prostate tumors that do not harbor deleterious mutations in these particular genes can similarly be deficient in HR, rendering them sensitive to HR-directed therapies.

To identify a more comprehensive set of prostate cancer cases with homologous recombination deficiency (HRD) including those cases that do not harbor mutations in known HR genes.

HRD levels can be estimated using various mutational signatures derived from next-generation sequencing data. We used this approach to determine whether prostate cancer cases display clear signs of HRD in somatic tumor biopsies. Whole genome (n=311) and whole exome sequencing data (n=498) of both primary and metastatic prostate adenocarcinomas (PRAD) were analyzed.

**Results:** Known BRCA-deficient samples showed robust signs of HR-deficiency associated mutational signatures. HRD-patterns were also detected in a subset of patients who did not harbor germline or somatic mutations in BRCA1/2 or other HR related genes. Patients with HRD signatures had a significantly worse prognosis than patients without signs of HRD.

**Conclusions:** These findings may expand the number of cases likely to respond to PARP-inhibitor treatment. Based on the HRD associated mutational signatures, 5-8 % of prostate cancer cases may be good candidates for PARP-inhibitor treatment (including those with BRCA1/2 mutations).

## 1 Introduction

Ovarian and breast cancer are often associated with mutations in BRCA1 and BRCA2, the key enzymes of a specific DNA repair pathway, homologous recombination (HR). Inactivating mutations in these genes often render such tumors homologous recombination deficient (HRD) that lead to sensitivity to PARP inhibitors, a novel class of cancer therapy, developed precisely for such tumors based on the principle of synthetic lethality [1]. Several PARP inhibitors have been approved over the past several years for the treatment of appropriately selected ovarian and breast cancer cases [2]. Since BRCA1 and BRCA2 mutations are also present in prostate adenocarcinoma, in up to 5-10% of metastatic prostate cancer cases [3], it was recently shown that PARP inhibitors were also effective in such prostate cancer cases [4].

HRD can be induced by germline or somatic BRCA1/2 genetic mutations; however, it can also be present in tumors with intact BRCA1/BRCA2 genes. The list of genes involved in homologous recombination is far from complete. For example, loss of CHD1 was only recently implicated as a possible mechanism leading to HRD in prostate cancer [5]. HRD can also be induced through epigenetic suppression of gene expression e.g. DNA methylation of BRCA1 or RAD51 [6]. Therefore, sequencing panels of HR related genes will likely miss a subset of HR deficient cases. Because tumors deficient in HRD are sensitive to PARPi/platinum, the consequence of not identifying tumors that harbor HRD is that treatment with these agents may be delayed or not given. Identification of such tumors is clinically relevant, because translational studies indicated that likely homologous recombination deficient tumors without BRCA1 or BRCA2 mutations may also benefit from homologous recombination deficiency associated therapy [7]. A particularly promising approach for *functionally* evaluating HRD focuses on specific DNA aberrations, or “DNA scars” that result from HRD. This approach is based on the principal that the normal function of homologous recombination ensures the error free repair of double strand breaks; therefore, in its absence, specific DNA scars accumulate in the genome that are the resultants of the error prone repair of double strand DNA breaks that occur due to e.g. replication stress. There are HRD induced DNA aberrations ranging from single nucleotide variation-based mutations to large scale, Mb sized genomic rearrangements [8]. These have been combined into predictive HRD markers both in the CLIA [9] and experimental [10] setting.

These methods have proven to be a robust indicator of the absence of BRCA1/BRCA2 function both in the genomic analysis of clinical samples and in direct induction experiments [9–11]. A recent study suggested that germline BRCA2 mutation-associated prostate cancer possess some of the previously described HRD associated mutational signatures in whole genome sequencing data [12].

In the current study, we investigated two related questions: 1) whether BRCA1 or BRCA2 mutations are consistently associated with the HRD induced mutational signatures in a large set of whole genome and whole exome sequencing data, and 2) whether there are prostate cancer cases with HRD-associated mutational signatures that are *not* associated with germline or somatic BRCA1 or BRCA2 mutations, which would suggest that there might be prostate cancer cases with sensitivity to PARP inhibitor treatment even in the absence of mutations in key HR genes.

## 2 Methods

### 2.1 Patients and cohorts

In this study we analyzed 311 whole genome sequenced samples from 240 cases from the following cohorts (Table 1). For the PRAD-CA, EOPC-DE, PRAD-UK cohorts processed data was used in our analyses. In case of the other 4 cohorts we worked with the BAM files. To evaluate the signs of HR-deficiency, further 498 whole exome sequencing samples were analyzed from the TCGA. The normal and tumor BAM files were downloaded via the GDC Data Portal.

**Table 1:** Summary of analyzed WGS cohorts.

| **Cohort** | **# patients** | **# samples** | **reference** |
| --- | --- | --- | --- |
| PRAD-CA | 110 | 110 | [30] |
| EOPC-DE | 39 | 61 | [31] |
| PRAD-UK | 28 | 77 | [32] |
| TCGA | 20 | 20 | [33] |
| DFCI | 19 | 19 | new data, phs000661.v1.p |
| CPDR | 14 | 14 | [34] |
| Decker | 10 | 10 | [12] |

### 2.2 Mutation, Copy number, and Structural Variant Calling PRAD-CA, EOPC-DE, PRAD-UK cohorts

For the PRAD-CA, EOPC-DE, PRAD-UK cohorts, germline mutations and structural variants were obtained from the DCC Data Portal. Somatic mutation status, allele-specific copy number and structural variant data were accessed from https://pcawg.xenahubs.net.

### 2.3 Mutation, Copy number, and Structural Variant Calling DFCI, CPDR, Decker, TCGA WGS and WES cohorts

In case of the other four cohorts, germline mutations were called with HaplotypeCaller, while somatic point-mutations and indels were called using Mutect2 (GATK 3.8). The high fidelity of the reported variants was ensured by the application of additional hard filters on top of the tools’ default ones. Allele-specific copy number profiles had been estimated by using Sequenza [13]. Structural Variants were called using BRASS (v6.0.0 - https://github.com/cancerit/BRASS). Further details are available in the Supplementary Notes.

### 2.4 Genotyping – BRCA-status

The mutations were annotated using InterVar [14]. Variants predicted as pathogenic or likely pathogenic were considered deleterious, while variants with unknown significance were marked differently. Copy number status of BRCA1/2 were based on Sequenza results.

### 2.5 Mutational Signatures

Somatic point-mutational signatures were determined with the deconstructSigs R package [15], by using the cosmic signatures as a mutational-process matrix. The extraction of rearrangement signatures was executed as described previously. [8]

### 2.6 Genomic scar scores

The calculation of the genomics scar scores (loss-of-heterozygosity: LOH, large scale transitions: LST and number of telomeric allelic imbalances: ntAI [16]) were determined using the scarHRD R package. [17]

### 2.7 HRDetect

Due to the lack of sufficient numbers of bona fide HR-deficient cases within the prostate cancer cohorts, and to the dissimilarities among the cases involved in their corresponding studies, a prostate-specific HRDetect model could not be created. Instead, the weights of the original, breast cancer-specific, whole genome-based HRDetect model [10] were used to calculate the HRDetect scores of the WGS prostate samples. The scores of the WES samples were calculated by using the weights of a whole exome specific model, that was trained on 560 artificial whole exomes. The predictors were log-transformed and standardized within all prostate cancer cases (n=311 for WGS, and n=498 samples for WES). Both the scores and sample attributes are available in Supplementary Table 1-2.

## 3 Results

### 3.1 Frequency of BRCA gene aberrations in prostate cancer in the whole genome sequenced cohorts

From the 311 samples (240 cases), 25 (18) had somatic or germline BRCA1/BRCA2 mutations.

Two patients (CPCG0234-F1, A17D-0095_CRUK_PC_0095), carried BRCA1 germline mutations. One of the patients also had an accompanying somatic aberration in BRCA1 (somatic mutation, LOH, respectively) while the other had a somatic structural variant aberration (BRCA1-SAFB translocation).

Five patients had BRCA2 germline mutations. Of these, two had additional BRCA2 aberrations (LOH, structural variant). Nine patients had a somatic BRCA2 mutation, out of which 3 also had a copy number loss.

Two samples had a deep deletion in BRCA2. (Supplementary Table 1.)

Both germline and somatic BRCA1/BRCA2 mutations may be present without additional loss of copy, resulting in retained homologous recombination capacity.

### 3.2 Genomic scars based HRD measures, the HRD score

Loss of function of BRCA1 or BRCA2 is associated with a range of distinct mutational signatures that include: 1) A single nucleotide variation based mutational signature (“COSMIC signature 3” or “BRCA signature” as labeled in the original publication [18]), 2) a short insertions/deletions based mutational profile, often dominated by deletions with microhomology, a sign of alternative repair mechanisms joining double strand breaks in the absence of homologous recombination [11] 3) large scale rearrangements such as non-clustered tandem duplications of a given size range (mainly associated with BRCA1 loss of function) or deletions in the range of 1-10kb (mainly associated with BRCA2 loss of function) [8].

Scoring systems have been devised to quantify the degree of HRD. The HRD-LOH [19] is the number of 15 Mb exceeding LOH regions which do not cover the whole chromosome, number of Large Scale Transitions (LST) [20] are defined as a chromosomal break between adjacent regions of at least 10 Mb, with a distance between them not larger than 3Mb, and Number of Telomeric Allelic Imbalances (ntAI) [21] is the number of AIs (unequal contribution of parental allele sequences) that extend to the telomeric end of a chromosome. The sum of these scores is referred to as HRD score, as in previous publications [9].

Most BRCA-deficient prostate cancer cases showed elevated levels of all these genomic scar scores both in the WGS (Figure 1A) and WES cohorts (Figure 2A, detailed results are presented in Supplementary Figures 1-5). There were, however, two of the biallelic BRCA-deficient cases that did not show signs of HR-deficiency. Furthermore, a subset of the BRCA1/2 wild type specimens had similarly high values to the known BRCA mutants.

**Figure 1:**
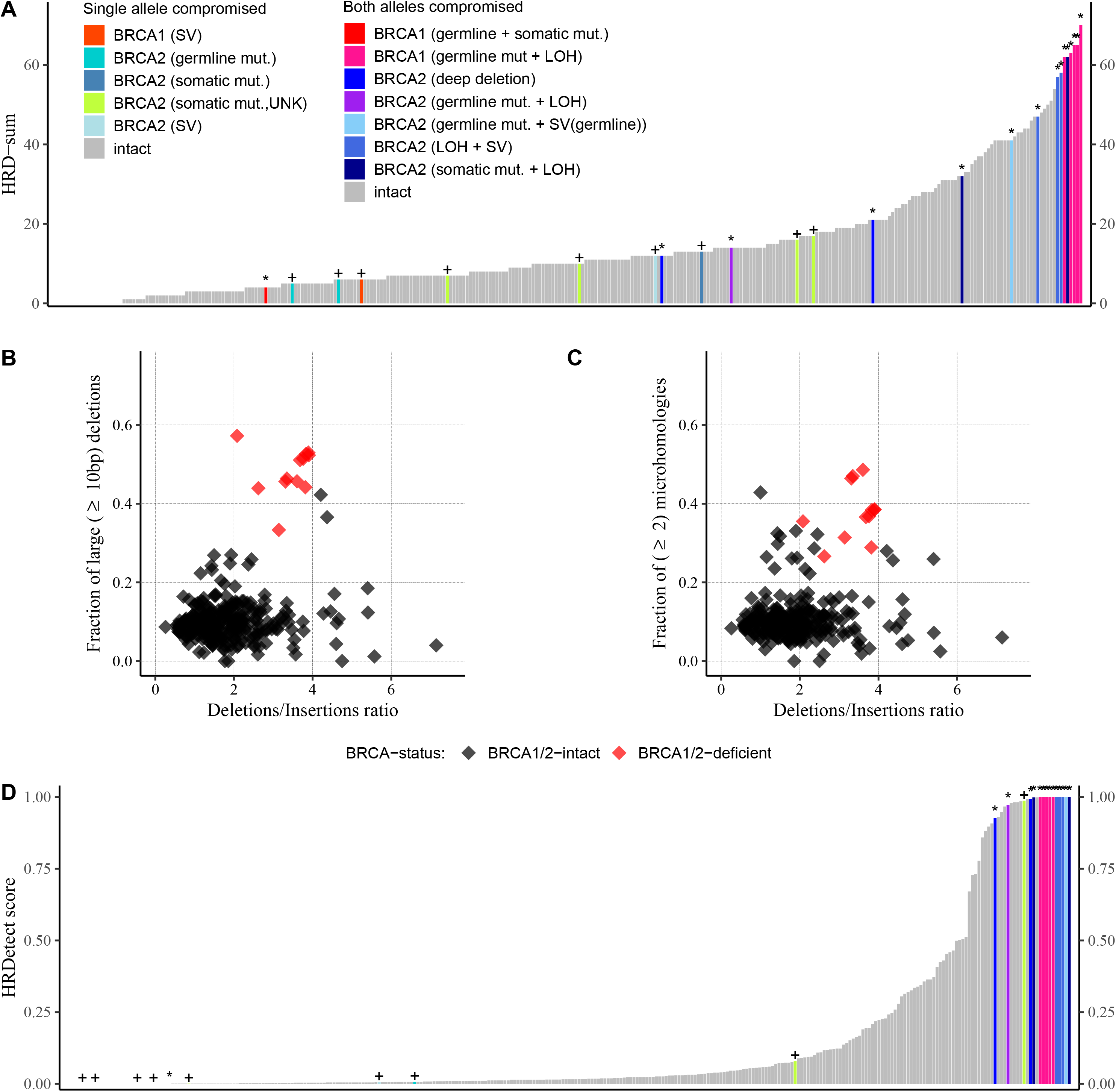
Summary of the HRD-related predictors in the whole genome datasets. **A:** HRD score: the sum of the three allele-specific CNV-derived genomic scars (HRD-LOH + LST + ntAI) **B:** Fraction of larger than 9 bp deletions versus deletions/insertions ratio **C**: Fraction of microhomology mediated deletions with larger or equal to 3 bp in length versus deletions/insertions ratio. **D:** HRDetect, the “+” signs if one allele is compromised, the “*” represent homozygous loss of BRCA1/2

**Figure 2:**
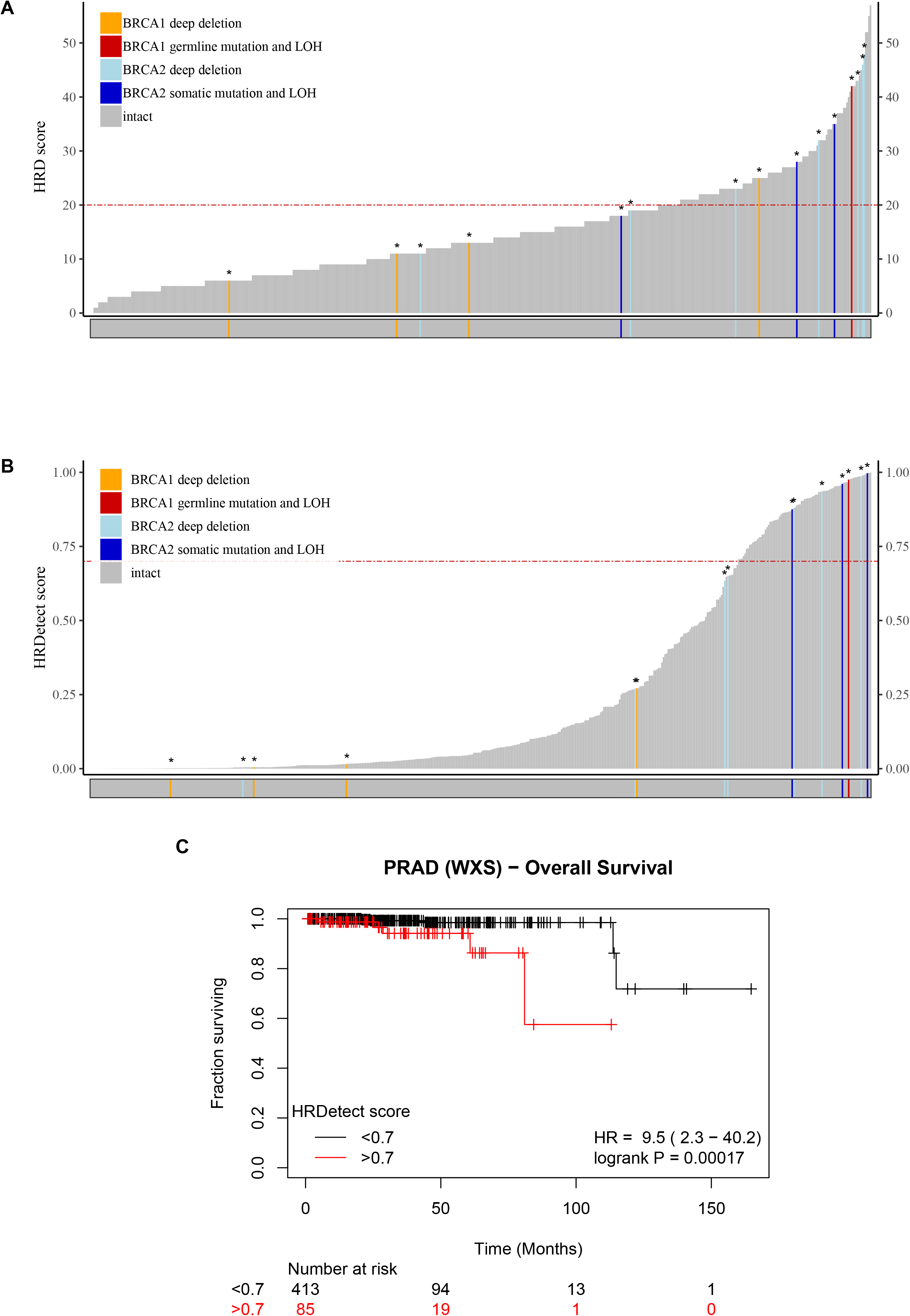
Summary of the HRD-related predictors in the whole exome datasets. **A:** HRD score per sample: the sum of the three allele-specific CNV-derived genomic scars (HRD-LOH + LST + ntAI) **B:** HRDetect the “+” signs if one allele is compromised, the “*” represent homozygous loss of BRCA1/2 **C:** Effect of HRDetect score on prognosis

### 3.3 Patterns of deletions and mutational signatures

The most robust signs of HR-deficiency, especially in a BRCA2−/−background, are the elevated Deletions/Insertions ratio, the relative increase in the number of deletions that are larger than 9 bp in length and the increased number of microhomology-mediated deletions (Figure 1B-C). Our analysis reinforces previous findings on larger cohorts [12].

BRCA1-mutations are known to be associated with a large contribution of Single Nucleotide Signature 3 [22]. Based on their signature composition, the DO51965_M1-M5 samples clearly demonstrate the signs of BRCA1-deficiency (Supplementary Figure 2).

Rearrangement Signature 3 and 5 has been associated with BRCA1 and BRCA2-deficiency, respectively. These signs were also detected in the investigated prostate cancer samples (Supplementary Figure 3).

### 3.4 HRDetect score

The HRDetect score was determined as described in the methods sections (section 2.7). Fourteen out of the 15 biallelic BRCA1/2 mutants among the WGS samples had larger than 0.7 HRDetect scores (Figure 1D), which is the current threshold of HR-deficient samples on whole genome sequences [10]. One BRCA1 mutant sample with both a germline and a somatic mutation acquired a low, close to zero HRDetect value, which can be explained with the assumption, that both of these mutations affected the same allele, leaving the other one intact, hence the true genotype might be +/−. Given that BRCA1/2 biallelic mutants are in effect HR-deficient, the HRDetect model had almost perfectly identified the known samples with HRD. These 14 samples however are not the only ones that have reached the higher ends of the HRDetect spectrum. 17 out of the 286 samples with a BRCA1/2 wild type background also reached above the 0.7 threshold, further strengthening the assumption, that even samples without mutations in the BRCA1/2 genes can exhibit the signs of HR-deficiency.

### 3.5 HR deficiency associated biomarkers in whole exome sequencing data

While whole exome sequences contain approximately 1% of the information that is available in their whole genome-derived counterparts, it has been previously shown that even that low information content might be enough to construct a reliable predictor for HR-deficiency. [23]

In case of the WES it is not possible to detect structural variants of BRCA1/2, thus lower number of identified BRCA1/2-mutants is expected. In the investigated cohort (n=498) four clearly BRCA2-deficient and one BRCA1-deficient cases were identified.

A modified, WES-based HRDetect score can be determined for these cases, which also demonstrates its ability to identify HRD cases. (Figure 2B)

Since a subset of the analyzed samples in the TCGA dataset had both WES and WGS data (n = 20), we performed a comparative analysis on the HRD-related attributes of the corresponding WES-WGS pairs (Supplementary Figure 6). We found, that the sum of the three genomic scars had a strong linear correlation (r = 0.79 +/− 0.14) between the sets, however this correlation was mainly driven by the number of large-scale transitions (r_LST_ = 0.92 +/− 0.09), while both the HRD-LOH and TAI scores exhibited weak positive correlation (r_HRD-LOH_ = 0.22 +/− 0.23 and r_TAI_ = 0.24 +/− 0.23 respectively). Similar weak correlations were observed between the WES and WGS-derived SNV signature 3 ratios (r = 0.21 +/−0.23) and microhomology-mediated deletion ratios (r = 0.23 +/− 0.23). (Figure 2B) HRDetect showed utility in case of WES, 7 out of 15 biallelic BRCA1/2 mutants had a score well above 0.7 (Figure 2 B). The HRDetect values for the other eight cases varied between 0 and 0.65. Although for whole exomes, this threshold is not well-established, it still indicates that the 78 out of 483 BRCA 1/2 wild type samples that have passed this value are also most plausibly HR deficient.

### 3.6 Mutational analysis of other HR-related genes

We investigated whether the high HRDetect scores in BRCA 1/2 wild type cases could be explained by the presence of biallelic aberration in other HR-related genes (such as BLM, PALB2, RAD50, RAD51 gene family, RAD52, RAD54B, and RAD54L). None of the WGS cases with higher than 0.7 HRDetect scores presented biallelic aberration in these genes. From the 55 WES cases that had higher than 0.7 HRDetect scores, 7 had biallelic BRCA1/2 mutations and additional 3 cases could be explained by biallelic mutations in PALB2, RAD51, or RAD54B. (Supplementary fig 10-11)

### 3.7 Intratumoral heterogeneity of HR-deficiency

Ten patients had multiple tumor samples sequenced. Eight of them had HRDetect values less than 0.3 in all of their specimens. For one patient, although the HRDetect scores varied between the tumor samples, they remained below 0.7 In case of EOPC-035 five out of the six tumor samples showed signs of HR-deficiency (Supplementary Figure 7).

### 3.8 Evolution of HR-deficiency

Ten patients from the investigated cohorts with metastatic prostate cancer had multiple metastatic sites sequenced. For two of the patients (A17 and A34) the primary prostate cancer and all of the metastatic sites showed signs of HR-deficiency and had higher than 0.9 HRDetect scores. For six of the patients (A10, A12, A21, A22, A24, A31) the primary tumor and all of the metastatic sites had low HRDetect scores, although in some samples (A21, A22, A31) slightly elevated HRDetect scores could be observed. Two patients (A29 and A32) showed heterogeneity in HR-deficiency, where although the primary prostate cancer was HR-competent, the metastases showed signs of HR-deficiency. (Supplementary Figure 8)

### 3.9 Effect of HRD on prognosis

In the WES-cohort, HRDetect-score higher than 0.7 was associated with worse overall survival (HR=9.5, p<0.01) (Figure 2C). This difference remained significant even after removing the germline BRCA1/2 mutant cases (Supplementary Figure 9). For the WGS cohorts, survival data was missing in the majority of cases, thus no significant difference could be detected.

## 4 Discussion

Personalized therapy of advanced metastatic prostate cancer has entered a new phase since the demonstration of the clinical efficacy of PARP inhibitors and platinum in cases with germline mutations in the DNA repair pathway, especially homologous recombination [4, 24]. However, some of the DNA damage checkpoint gene germline mutants do not benefit from PARP inhibitor therapy [25] and some cases without such germline mutations are sensitive to platinum-based therapy [24, 26]. Therefore, optimal use of these therapeutic approaches will require more accurate identification of HR deficient cases. A similar diagnostic problem was addressed in the case of breast and ovarian cancer using HR deficiency induced mutational signatures [10, 27]. We decided to apply the same approach here. First, we demonstrated that bona fide, key HR gene (BRCA1/2) deficient cases show robust signs of HR deficiency-induced mutational signatures. Second, we also found that there is a significant number of cases without germline or somatic mutations in these genes that also demonstrate robust signs of HR deficiency-induced mutational signatures. This strongly suggests that there is an additional, identifiable set of metastatic prostate cancer cases that will likely benefit from PARP inhibitor-based therapy. In support of assumption, a recent clinical report found at least one case without germline mutations of HR genes with exceptionally good response to platinum based therapy showing HRD+ score based on WES analysis [26]. A clinical trial, prioritizing patients based on such molecular signatures will be needed to further validate this hypothesis.

We could explain the presence of HRD induced mutational signatures with mutations in other HR related genes, such as PALB2, RAD51, or RAD54B, in a few cases. HRD can also be caused by the low expression of HR related genes. However, detecting correlations between low expression of HR genes and HRD induced mutational signatures is severely limited by factors such as normal tissue contamination. For example, significant expression deficiency or LOH of the BRCA1 or BRCA2 genes can often be masked by the presence of these genes in the normal cells in the tumor biopsy.

Mutational signatures can be most accurately determined from whole genome sequencing (WGS) data since, for example, large scale rearrangements can be detected only in such data. Since WGS data are not always available, we investigated whether whole exome sequencing could also serve as a reliable detector of HR deficiency. WES data was able to provide a robust, though possibly less sensitive, measure of HR deficiency of clinical cases. Despite the wide availability of next generation sequencing (NGS), no CLIA certified WGS or WES based methods are available for determining HR deficiency in the diagnostic setting. Currently only targeted-sequencing based estimation of the genomic scar scores are available in CLIA-certified setting such as the myChoice assay (Myriad Genetics) for breast and ovarian cancer. With this method, tumors are considered HRD+ if they have a high myChoice HRD score (≥42) or a tumor BRCA1/2 mutation and HRD- if they have a low myChoice HRD score (<42) and wild-type BRCA1/2. [9] Unfortunately our analysis didn’t have the sufficient number of BRCA-mutant cases to determine a prostate cancer-specific cutoff value, as in the case of ovarian cancer [9]. Nevertheless, the BRCA-deficient cases have clearly higher genomic scar scores both in the WES and the WGS cohorts. (Supplementary Figure 4-5). A significantly higher number of cases will be needed to determine the clinically useful threshold value.

It has been previously demonstrated that [28] that metastasis to metastasis spread of prostate tumor clones is common. This may explain the small heterogeneity seen in HR-status of multiple metastases.

Prostate cancer with BRCA2 germline mutation is known to be associated with poorer prognosis [29]. In our analyses we were not able to demonstrate this effect due to low number BRCA-mutant cases. However, we found that tumors with higher than 0.7 HRDetect-score have worse overall survival, which would also warrant their expedited involvement in targeted, PARP inhibitor based trials.

## 5 Conclusions

In summary, we show that the presence of BRCA1/2 mutations is associated with the presence of HR-deficiency associated mutational signatures in prostate adenocarcinoma. HRD-patterns were also detected in a subset of patients who did not harbor germline or somatic mutations in BRCA1/2 or other HR related genes. This likely defines a subset of prostate cancer patients that will likely benefit from PARP inhibitor or platinum-based therapy.

## Acknowledgment

This work was supported by the Research and Technology Innovation Fund (KTIA_NAP_13-2014-0021 to Z.S.); Breast Cancer Research Foundation (BCRF-17-156 to Z.S.) and the Novo Nordisk Foundation Interdisciplinary Synergy Programme Grant (NNF15OC0016584 to Z.S.), Department of Defense through the Prostate Cancer Research Program (award number is W81XWH-18-2-0056) to Z.S. and M.F. The Danish Cancer Society grant (R90-A6213 to MK). Z.S. and J.B. was supported by Velux Foundation 00018310 grant. The results shown here are based upon data generated by the TCGA Research Network: http://cancergenome.nih.gov/ and the International Cancer Genome Consortium (ICGC): https://icgc.org/.

## Availability of data and material

Data analysis was carried out using R, version 3.5.0. Sequencing data that support the findings of this study have been deposited in the dbGaP with the accession code phs000661.v1.p and the previous published datasets repositories are listed in the supplementary material.

## Authors contribution

ZsS and MD designed the analysis and performed all computational analysis and participated in preparation of the manuscript. MK, JB, MP, VT, SS, OR, IC, and MF participated in designing the analysis and prepared the manuscript. ZS designed the analysis, had supervisory role and prepared the manuscript.

