## Supplementary notes for "Genomic aberration based molecular signatures efficiently characterize homologous recombination deficiency in prostate cancer"

---

#### Supplementary Material

---

*Written by:*

Zsofia Sztupinszki, Miklos Diossy, Marcin Krzystanek, Judit Borcsok, Mark Pomerantz, Viktoria Tisza, Sandor Spisak, Orsolya Rusz, Istvan Csabai, Matthew Freedman, Zoltan Szallasi

### Contents

1 Supplementary Methods 1
2 Genomic scar scores 1
3 Mutational signatures 5
4 SV signatures 8
5 WES HRD predictors 11
6 Genomic scar scores in WGS 12
7 WES vs WGS predictors 13
8 HRD-ITH 14
9 HRDetect-ITH-Mets 15
10 HRD-prognosis 18
11 Genotyping HR-related genes - WGS 19
12 Genotyping HR-related genes - WES 19

### 1 Supplementary Methods

**Mutation, Copy number, and Structural Variant Calling**

**PRAD-CA, EOPC-DE, PRAD-UK cohorts:**
For the PRAD-CA, EOPC-DE, PRAD-UK cohorts germline mutations and structural variants were obtained from the DCC data portal (SNPs, InDels and complex short variants were called using GATK HaplotypeCaller, FreeBayes and RTG, SVs were called using Delly and the TRAFIC mobile element insertion caller.) Somatic mutation status (PCAWG-1 Consensus Pass-Only Somatic Whole Genome Mutation Calls, union callset of the passing variants from the three core callers (Sanger, Broad, DKFZ-EMBL), plus MuSE 1.0, and filtering out calls made by only one calle), allele-specific copynumber and structural variant (PCAWG6 Structural Variant merge set v 1.6.Merge is based on SV calls from BRASS (Sanger), DELLY (EMBL-DKFZ), dRanger and SNOWMAN (BROAD),. The data were accessed from https://pcawg.xenahubs.net

**DFCI, CPDR, Decker, TCGA WGS and WES cohorts:**
In case of the other four cohorts, germline mutations were called with HaplotypeCaller, and somatic point-mutations and indels had been called using Mutect2 (GATK 3.8). The high fidelity of the reported variants was ensured by the application of additional hard filters on top of the tools' default ones. Only those germline variants were considered that had a coverage of at least 15 and a variant quality of at least 20 (PHRED). The minimum coverage of the somatic variants was set to 15 in the normal and 20 in the tumor, and they also had to have a tumor LOD (logarithm of odds) of at least 6, and a normal LOD of 4. The minimum tumor allelic frequency was set to 0.05. Allele-specific copy number (CN) profiles had been estimated by using sequenza[1]. The grid-approximated posterior space was defined in the ploidy range [1,5] (0.1 bin size) and cellularity range [0,1] (0.01 bin size). When the maximum a posteriori estimate resulted in an unlikely CN profile (e.g. an unrealistically high ploidy coupled with a low cellularity), an alternative mode of the posterior was selected manually. Structural Variants were called using BRASS (v6.0.0 - https://github.com/cancerit/BRASS)

**MSI-status**

The MSI-status of the samples were determined with MSISeq[2], and all of the samples proved to be MSS. Also, the contribution of Signature 6 was negligible in all of the cases.

### 2 Supplementary Figure 1, Genomic scar scores

Legend for the colors of the x-axis for the following plots:

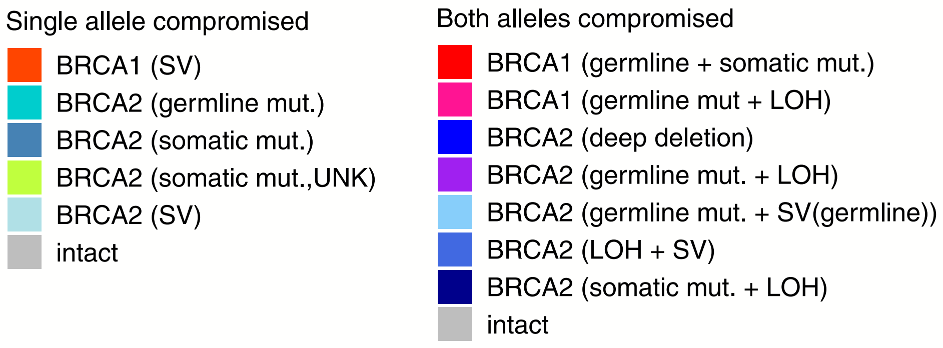

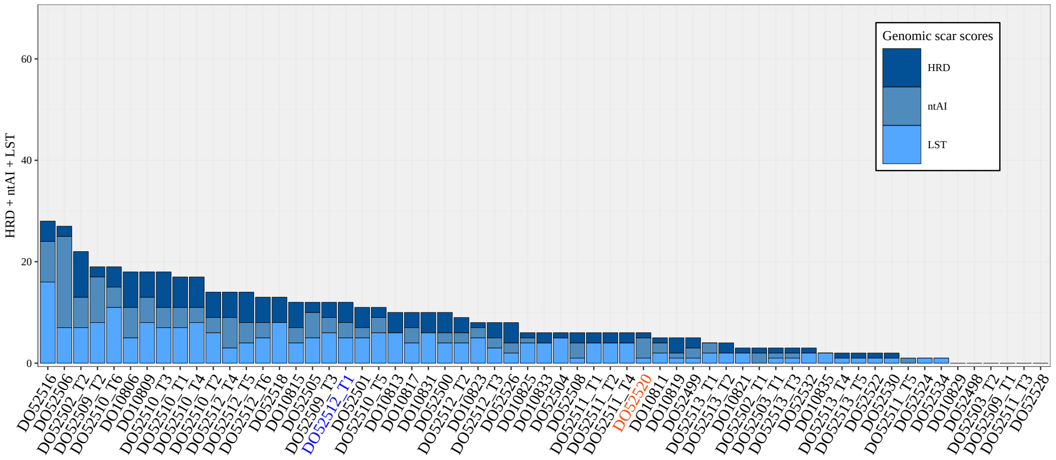


**(A)** EOPC-DE cohort

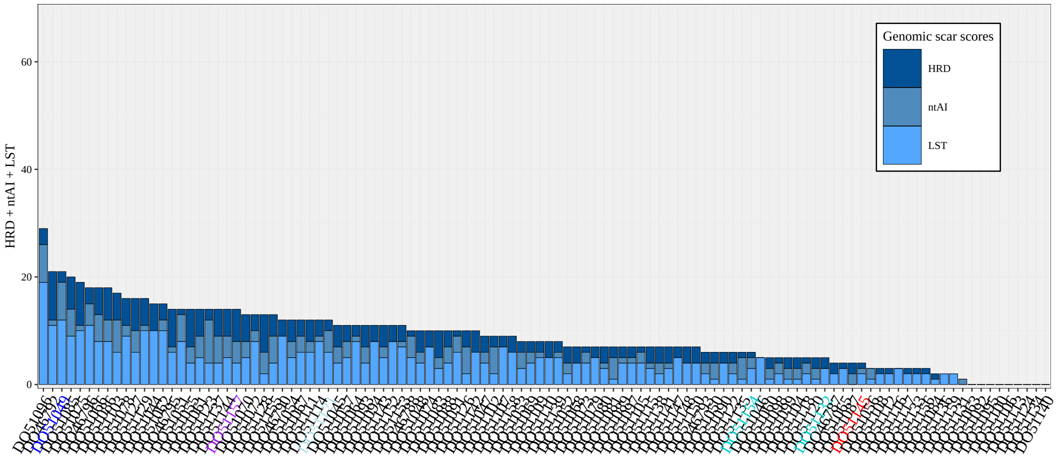


**(B)** PRAD-CA cohort

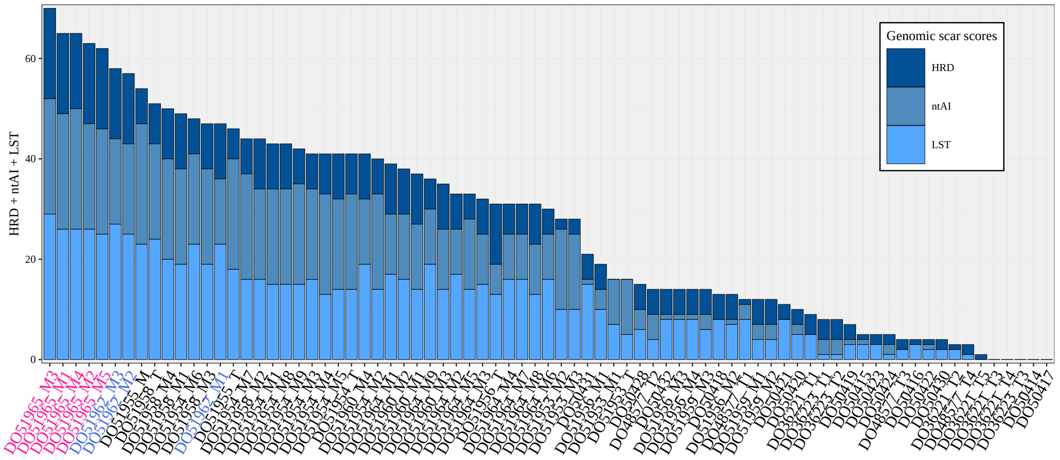


**(C)** PRAD-UK cohort

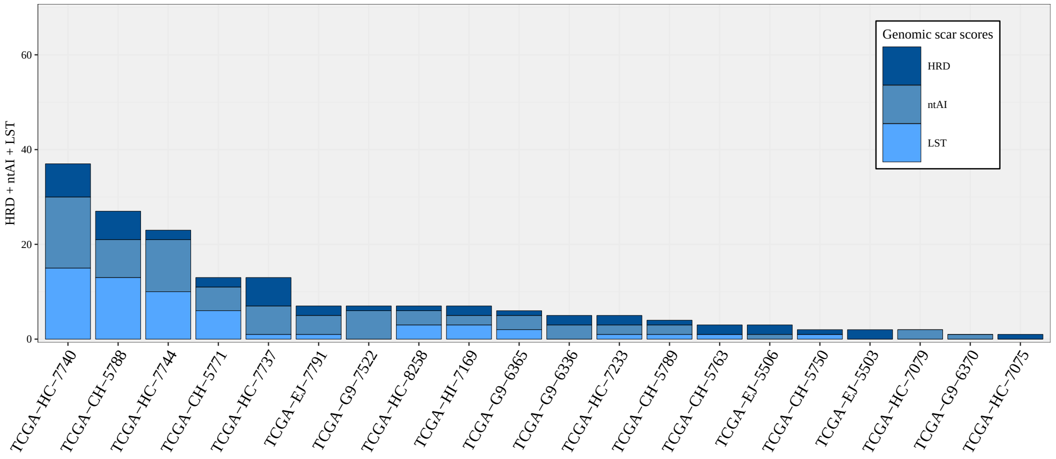


**(D)** TCGA cohort

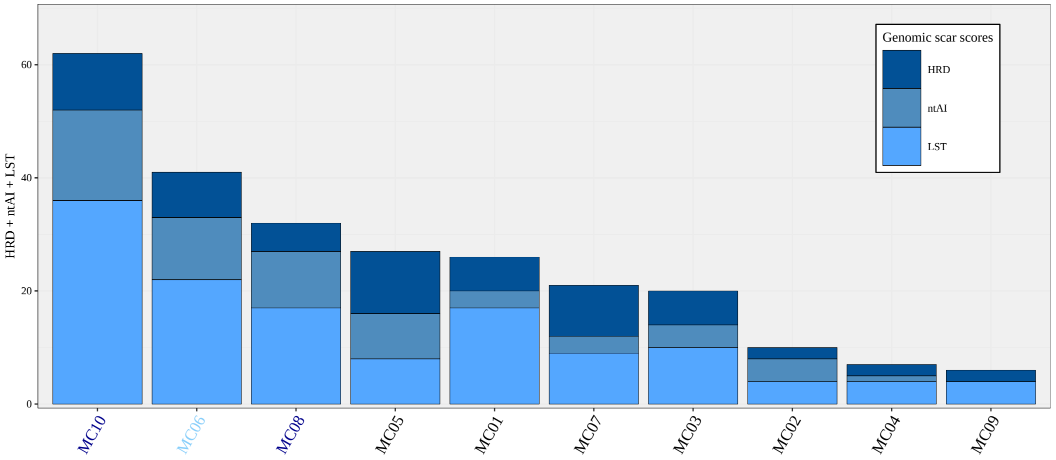


**(E)** Decker cohort

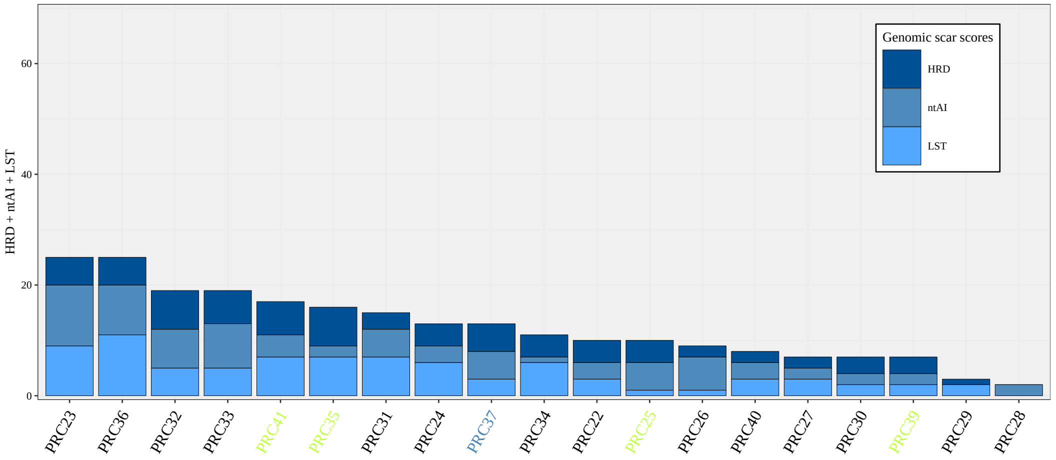


**(F)** DFCI cohort

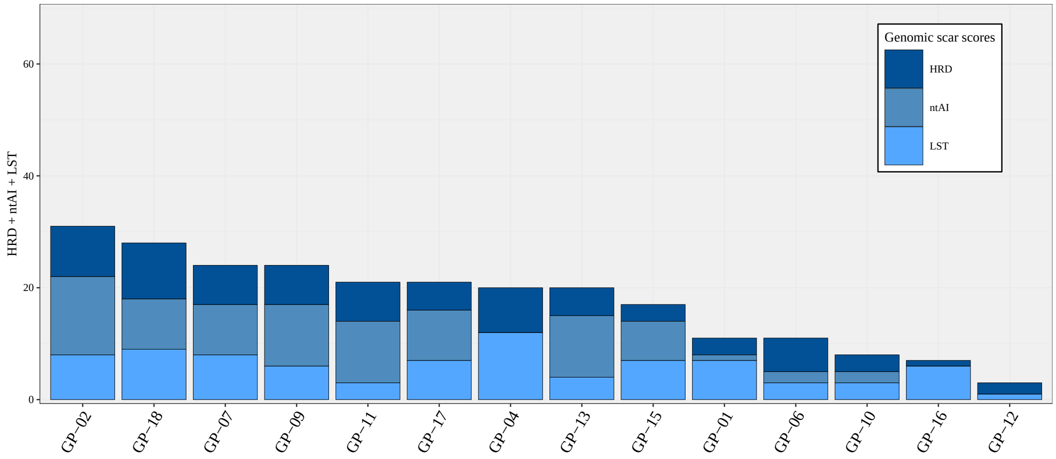


**(G)** CPDR cohort

**Suppl.Fig. 1:** The genomic scar scores (HRD-LOH, nTAI, LST) per sample grouped by the cohorts.

### 3 Supplementary Figure 2, Relative contribution of Mutational Signatures

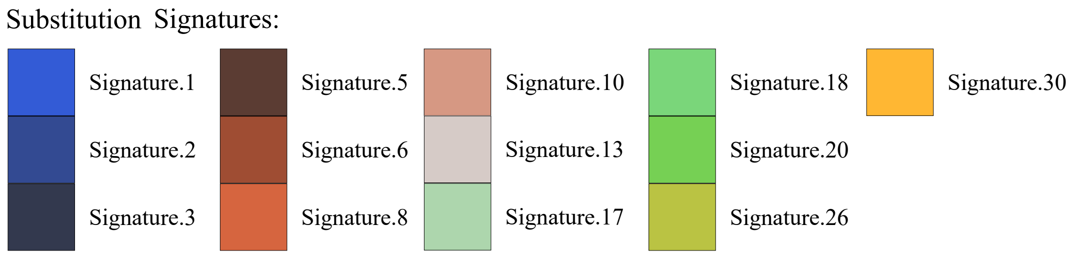


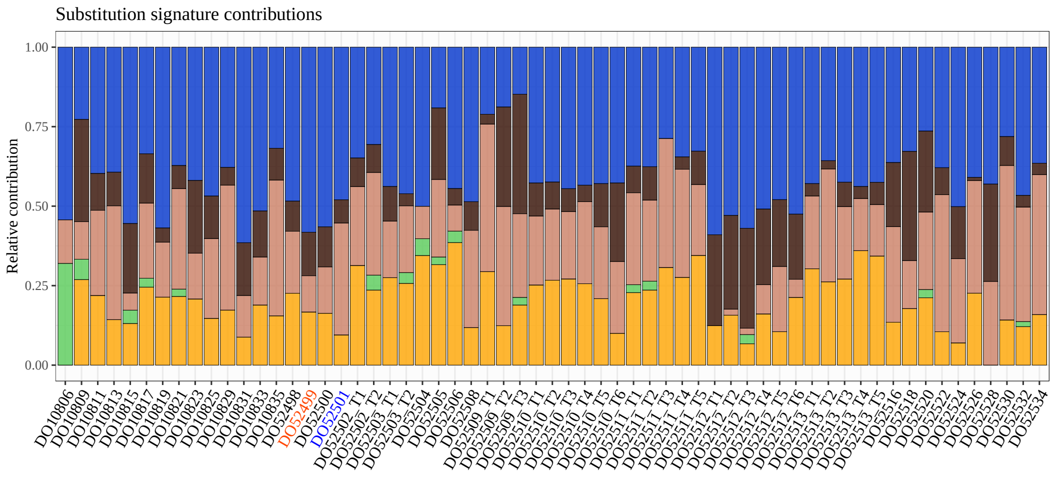


**(A)** EOPC-DE cohort

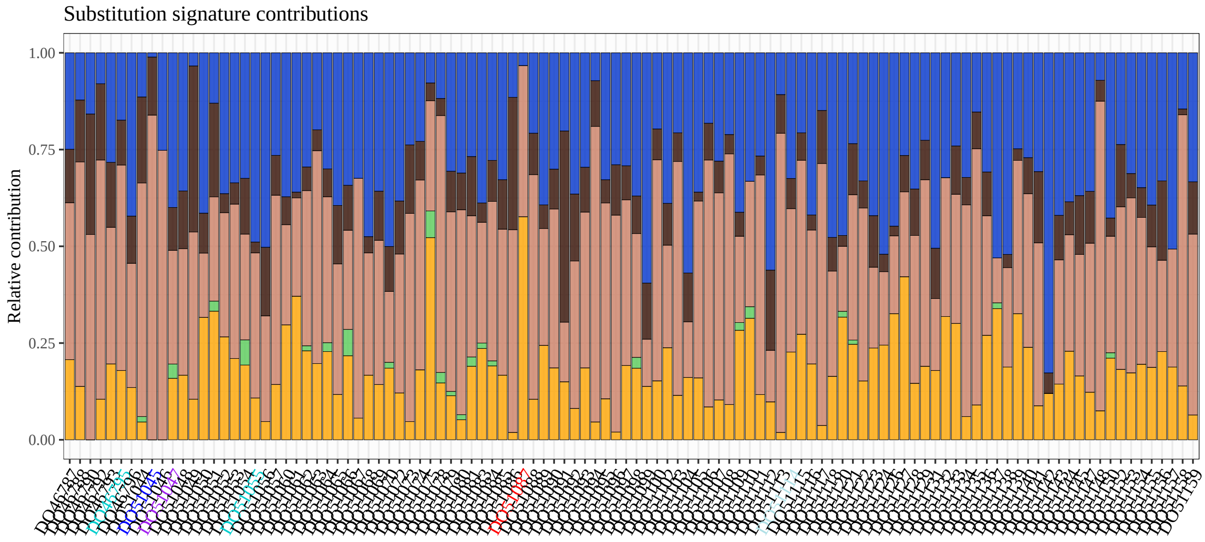


**(B)** PRAD-CA cohort

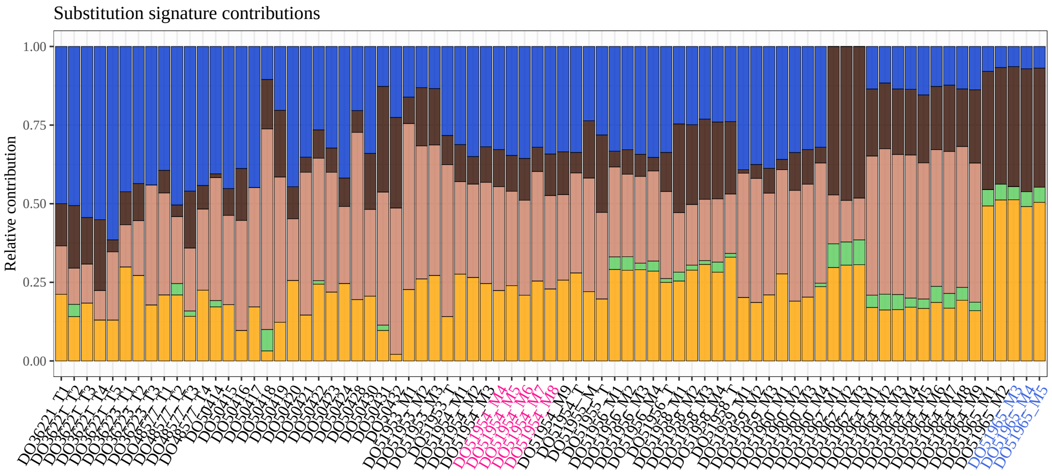


**(C)** PRAD-UK cohort

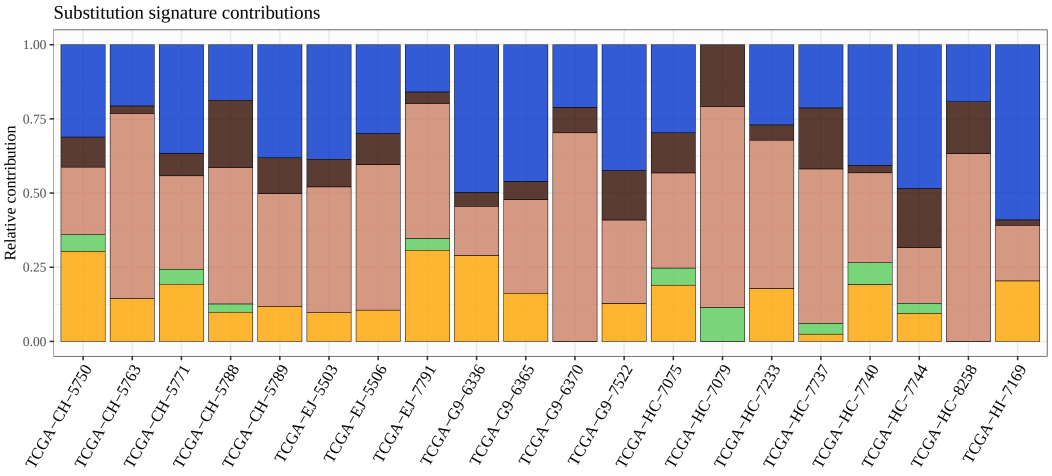


**(D)** TCGA cohort

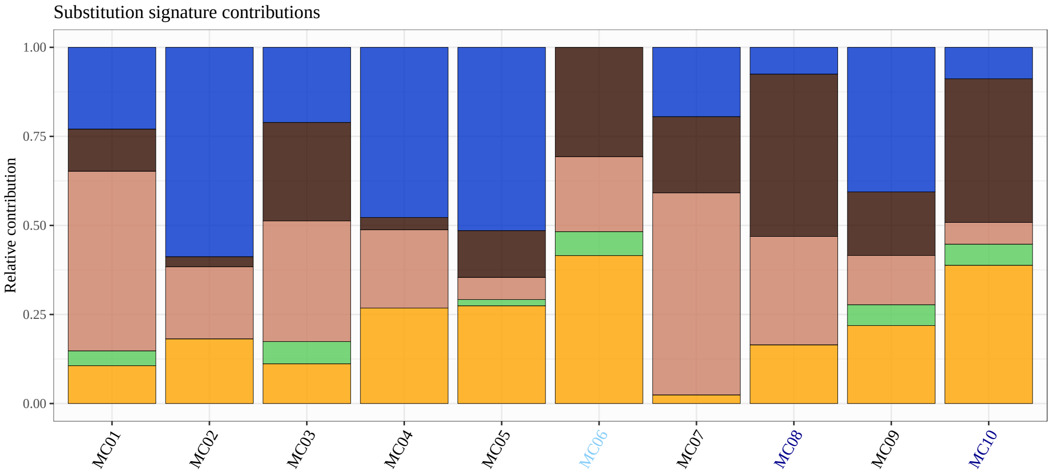


**(E)** Decker cohort

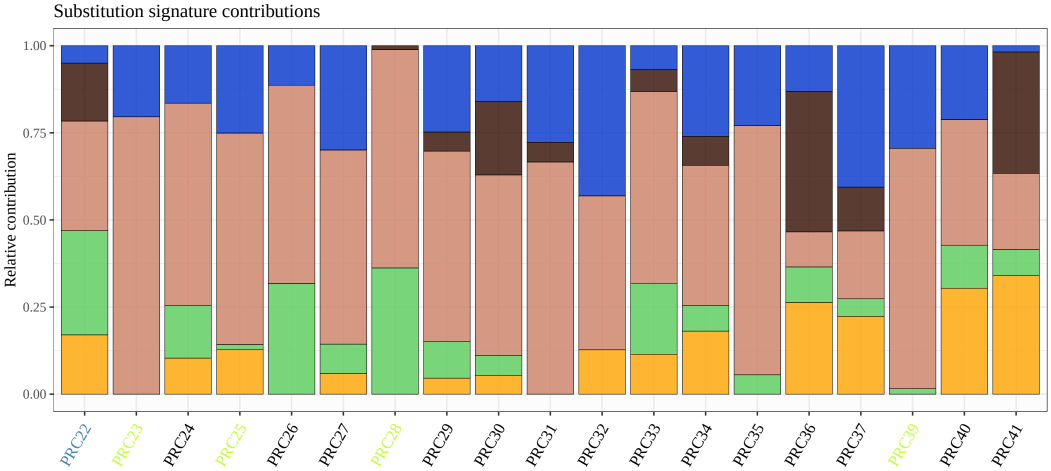


**(F)** DFCI cohort

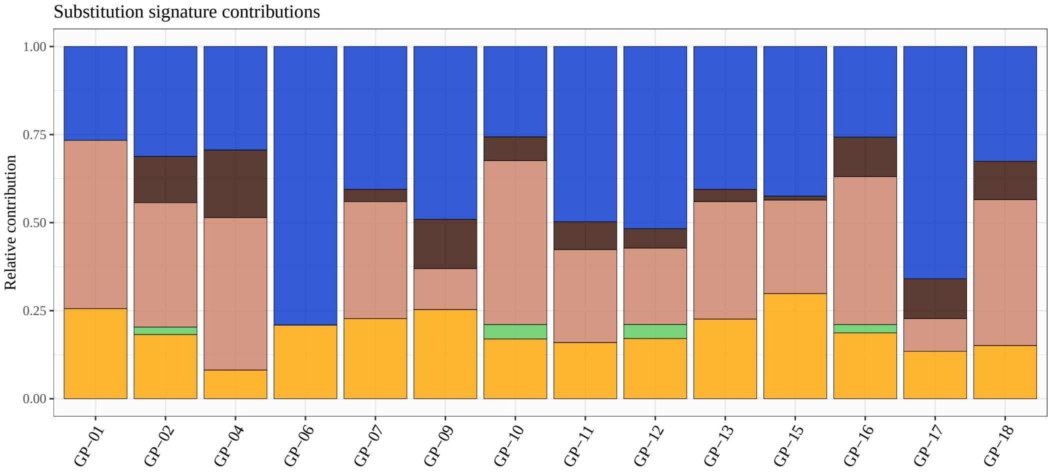


**(G)** CPDR cohort

**Suppl.Fig. 2:** Contribution of substitutional signatures per patient

### 4 Supplementary Figure 3, Relative contribution of Rearrangement Signatures

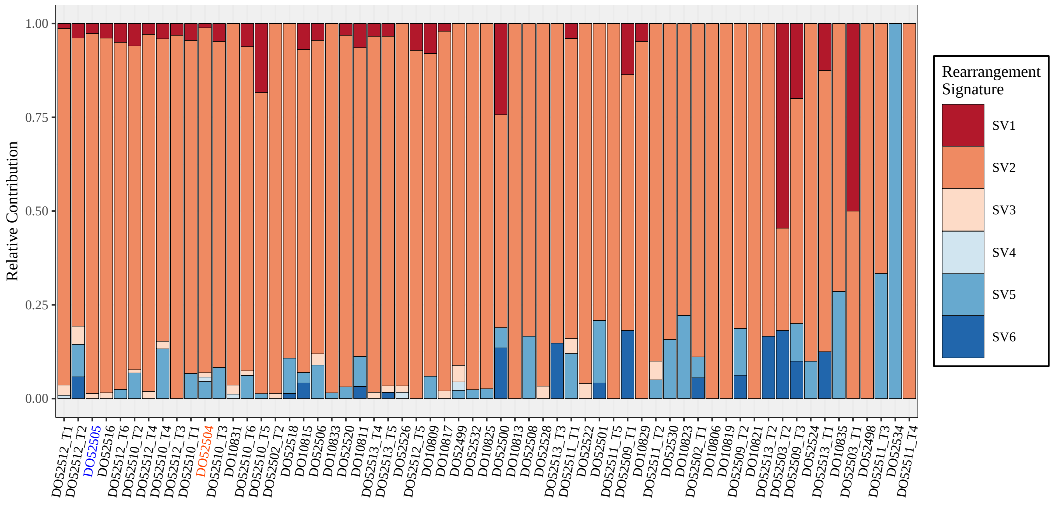


**(A)** EOPC-DE cohort

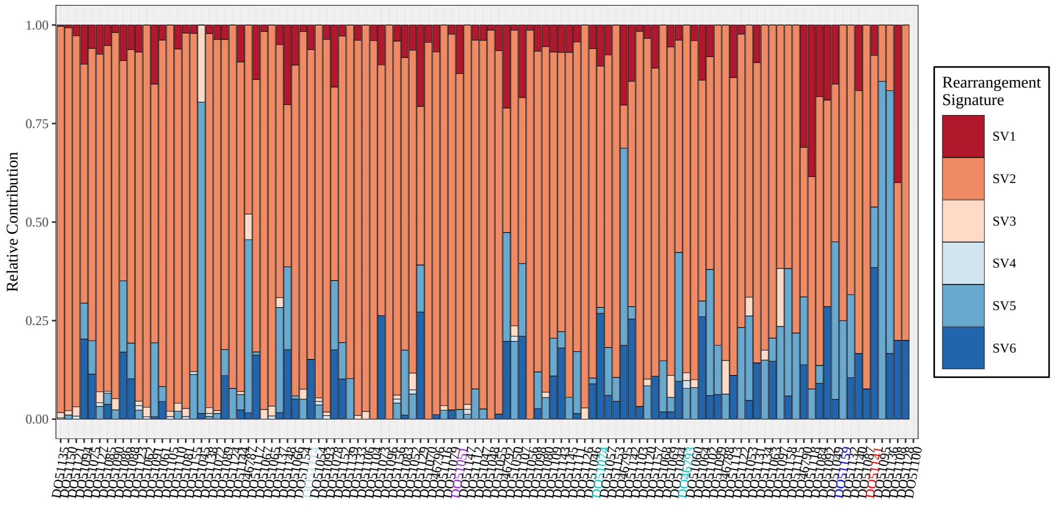


**(B)** PRAD-CA cohort

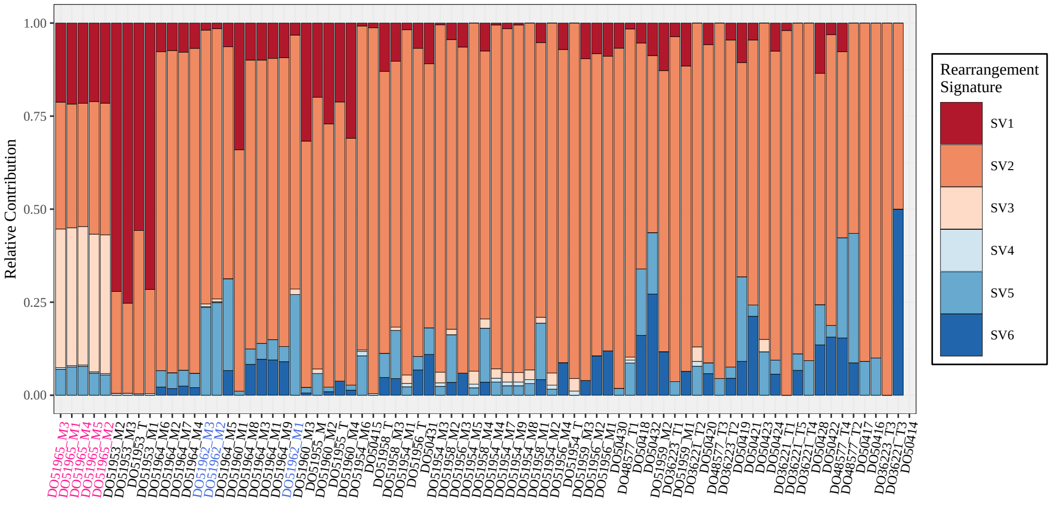


**(C)** PRAD-UK cohort

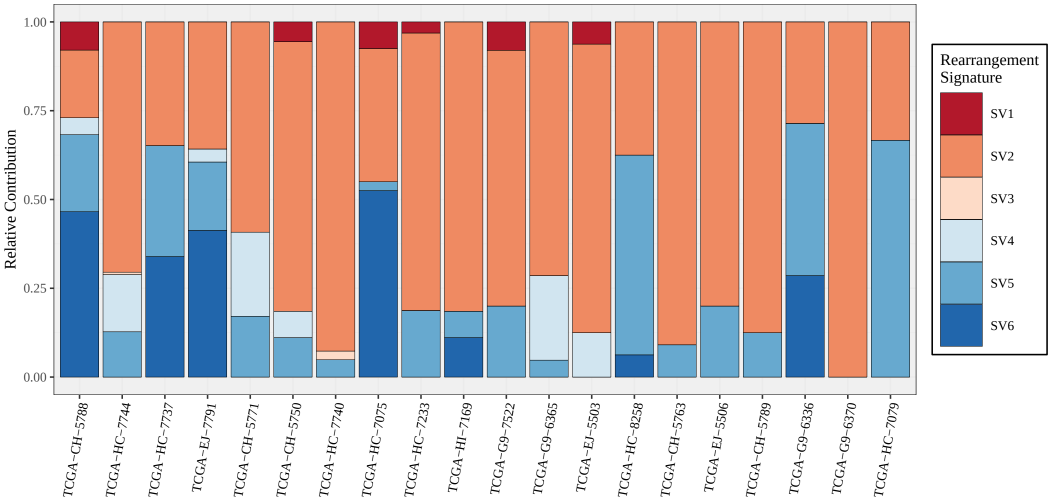


**(D)** TCGA cohort

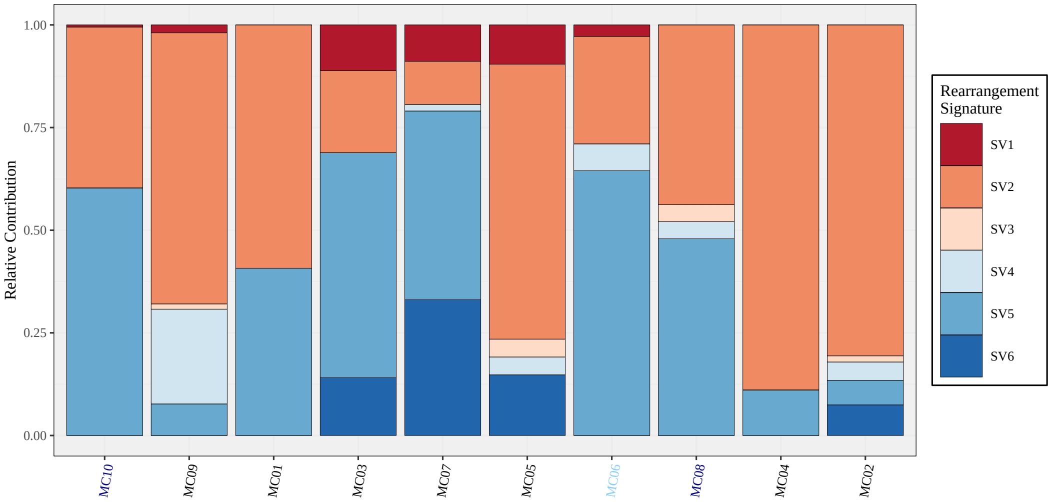


**(E)** Decker cohort

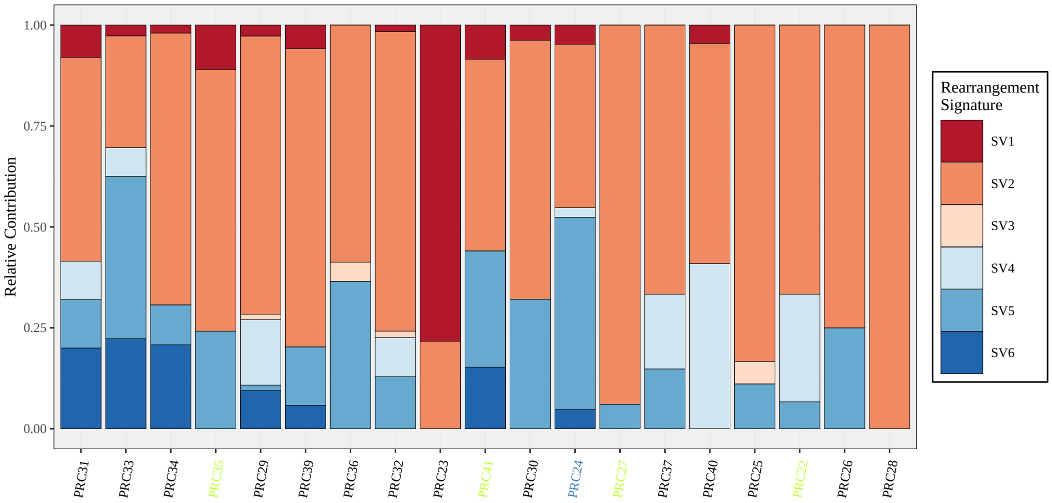


**(F)** DFCI cohort

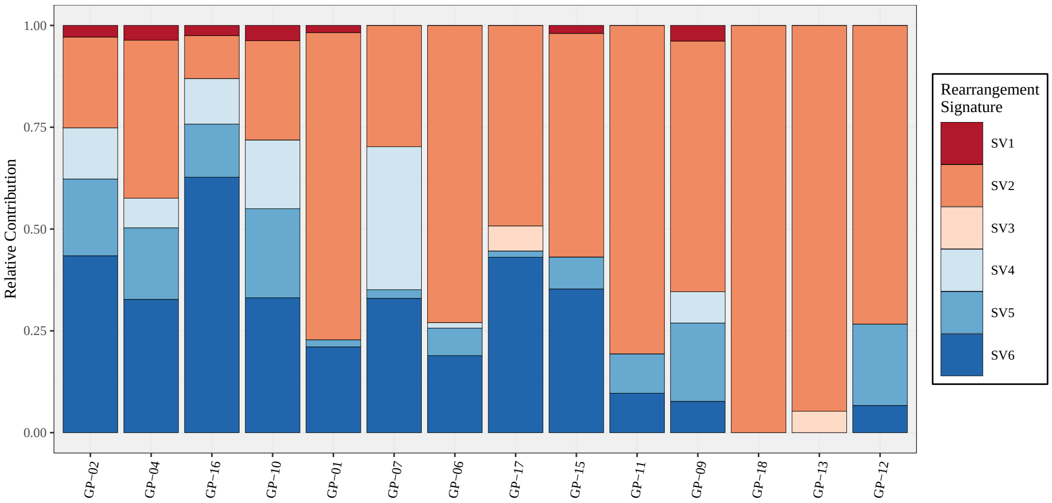


**(G)** CPDR cohort

**Suppl.Fig. 3:** Relative contribution of Rearrangement Signatures per sample grouped by the cohorts.

### 5 Supplementary Figure 4, HRD-related genomic features extracted from the whole exomes

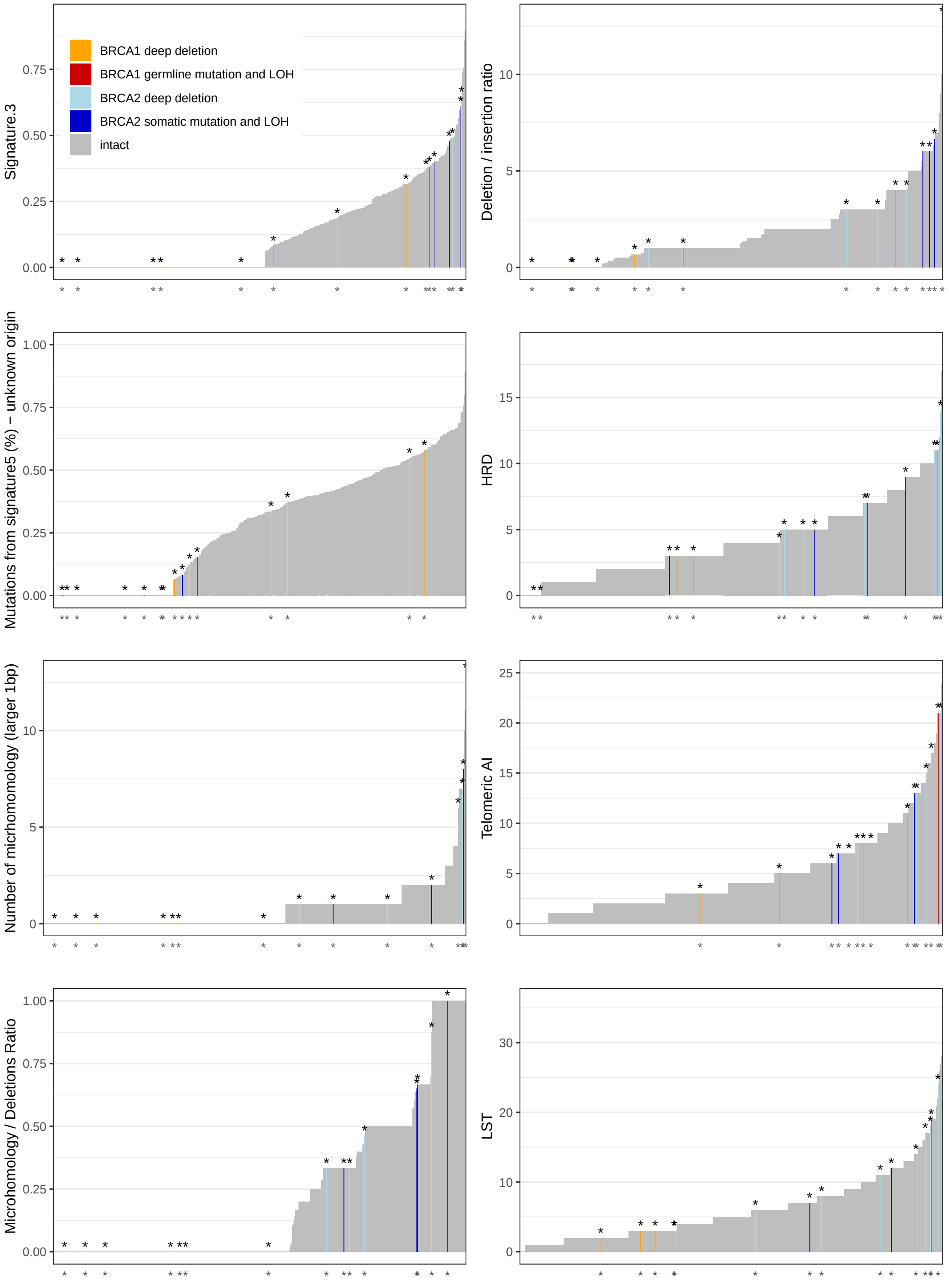


**Suppl.Fig. 4:** *Distribution of genomic scar scores (HRD-LOH, Telomeric Allelic Imbalance, Large-scale Transitions), homologous-recombination deficiency related mutational signatures (Signature 3, 5), number of micrhomomology-mediated deletions, microhomology / deletions ratio, deletion / insertion ratio and BRCA1/2-status in whole exome sequnced prostate adenocarcinoma samples (n=498).*

### 6 Supplementary Figure 5, Genomic scar scores in WGS

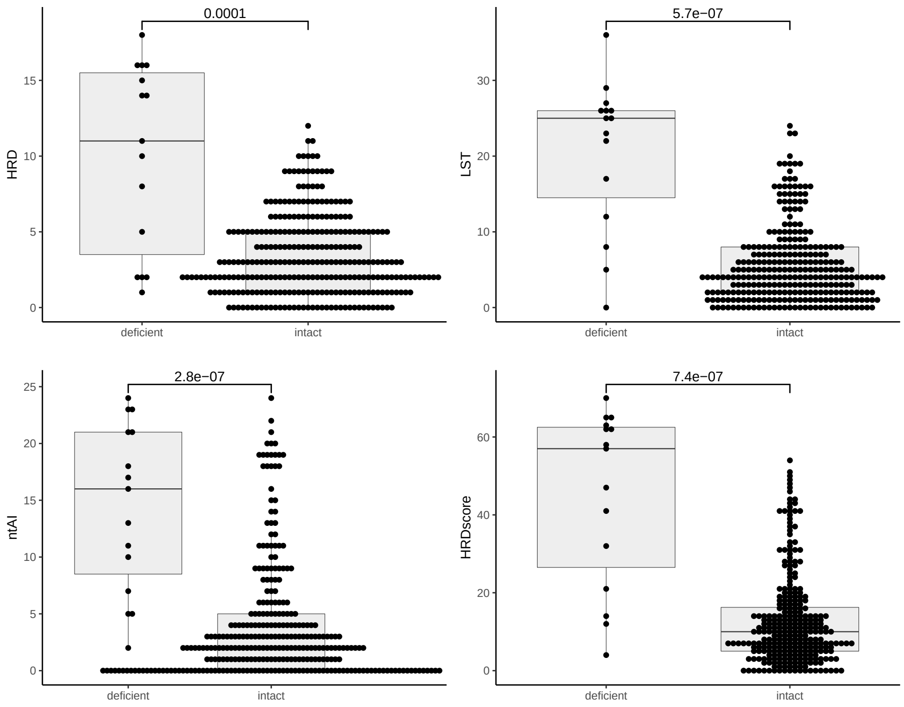


**Suppl.Fig. 5:** Genomic scar scores in WGS

### 7 Supplementary Figure 6, Correlations between the WES and WGS HRD-predictors

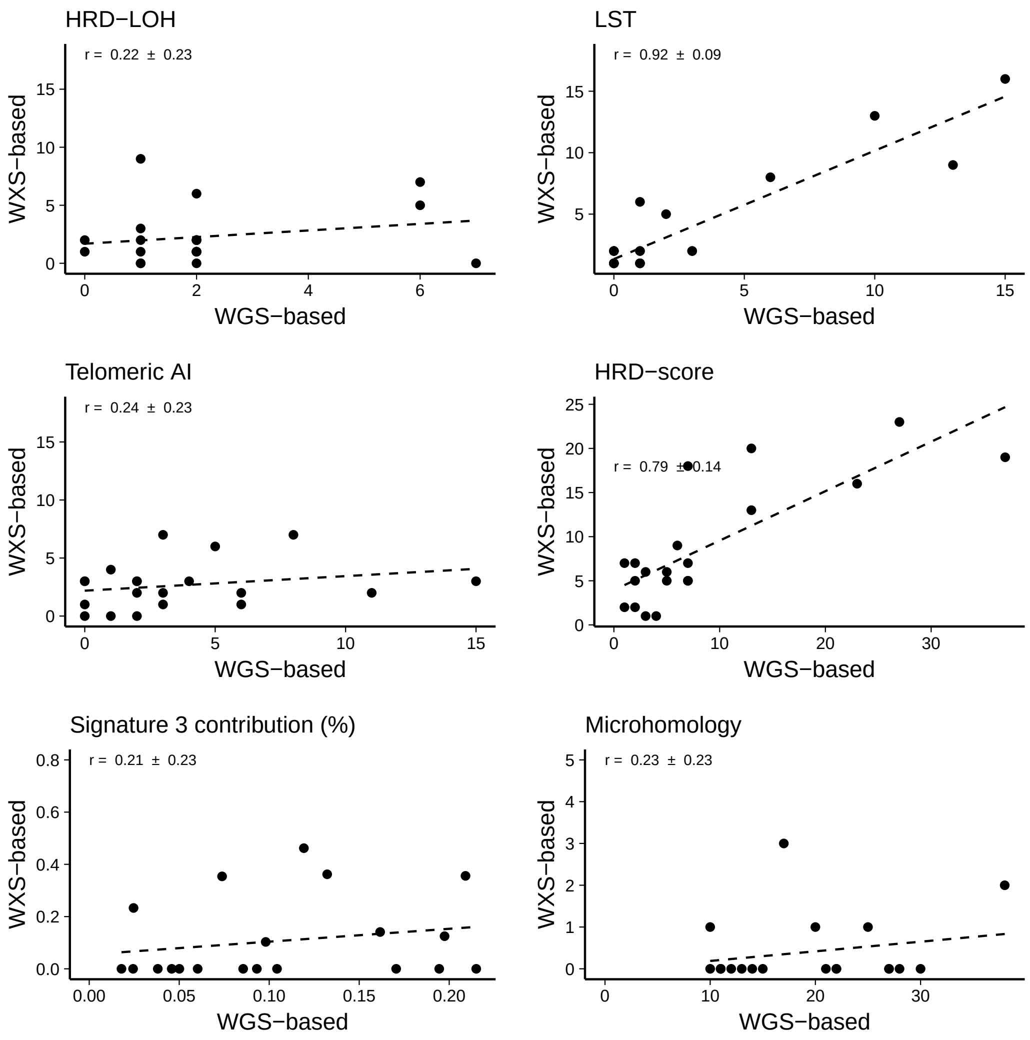


**Suppl.Fig. 6:** *Correlation of the three main components (number of HRD-LOH events, number of microhomology-mediated deletions, and contribution of Signature 3 to the mutational profile) of HRDetect between paired whole exome and whole genome sequenced samples*

### 8 Supplementary Figure 7, Intratumor heterogeneity of HR-deficiency

For 10 patients multiple tumor samples were available (from 2 to 6 samples per case). In some cases the WGS-based HRDetect-score showed mild intratumor heterogeneity.

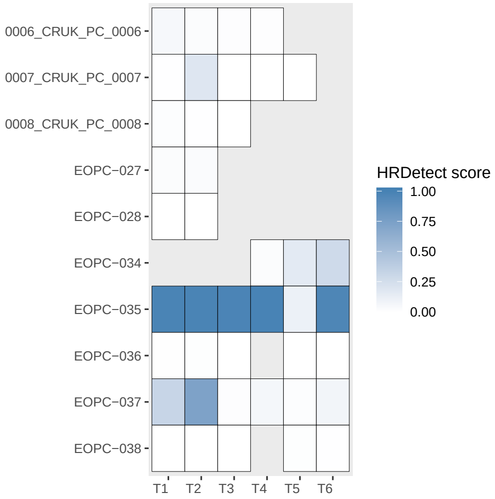


**Suppl.Fig. 7:** HRDetect-score in multiple samples from the same patients

### 9 Supplementary Figure 8, Heterogeneity of HR-deficiency between metastases

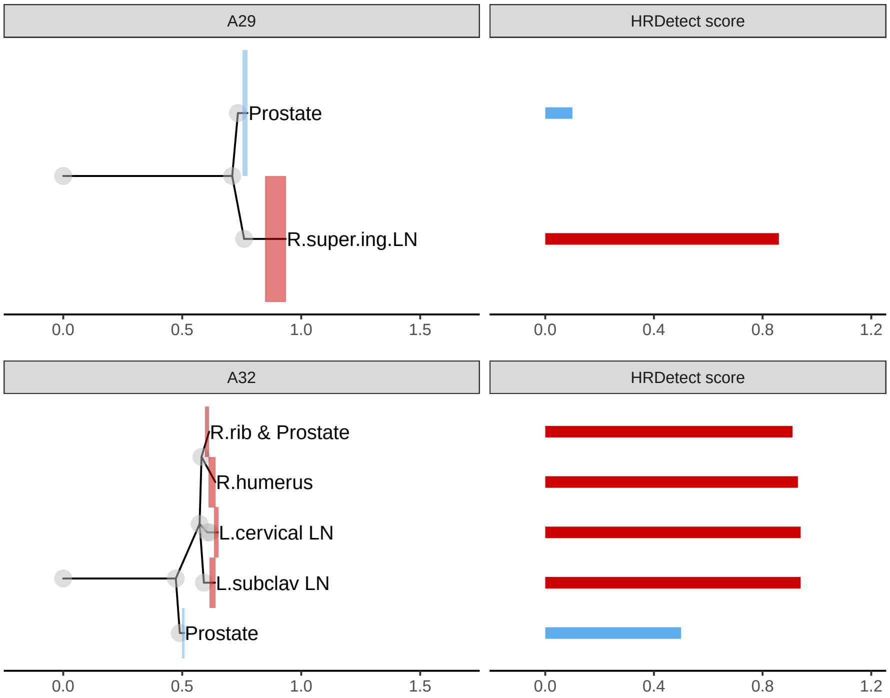


**(A)**

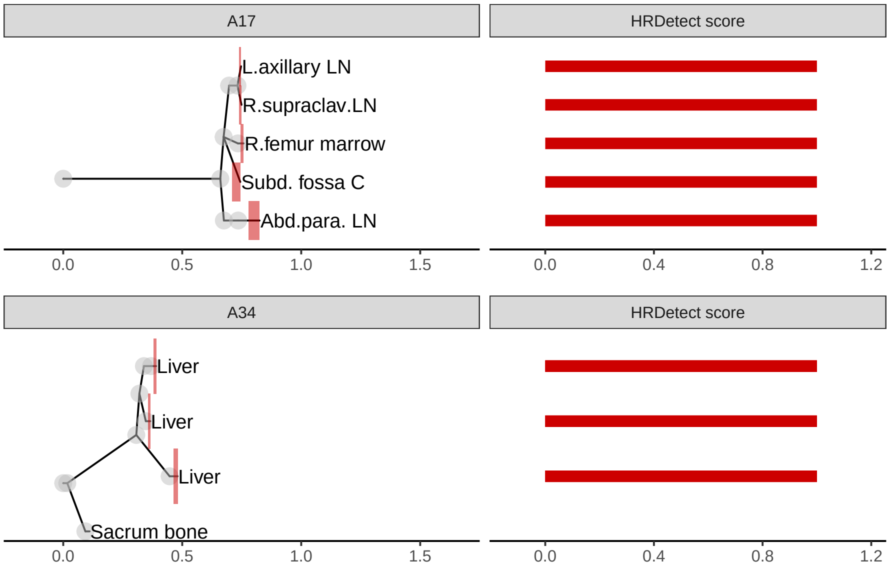


**(B)**

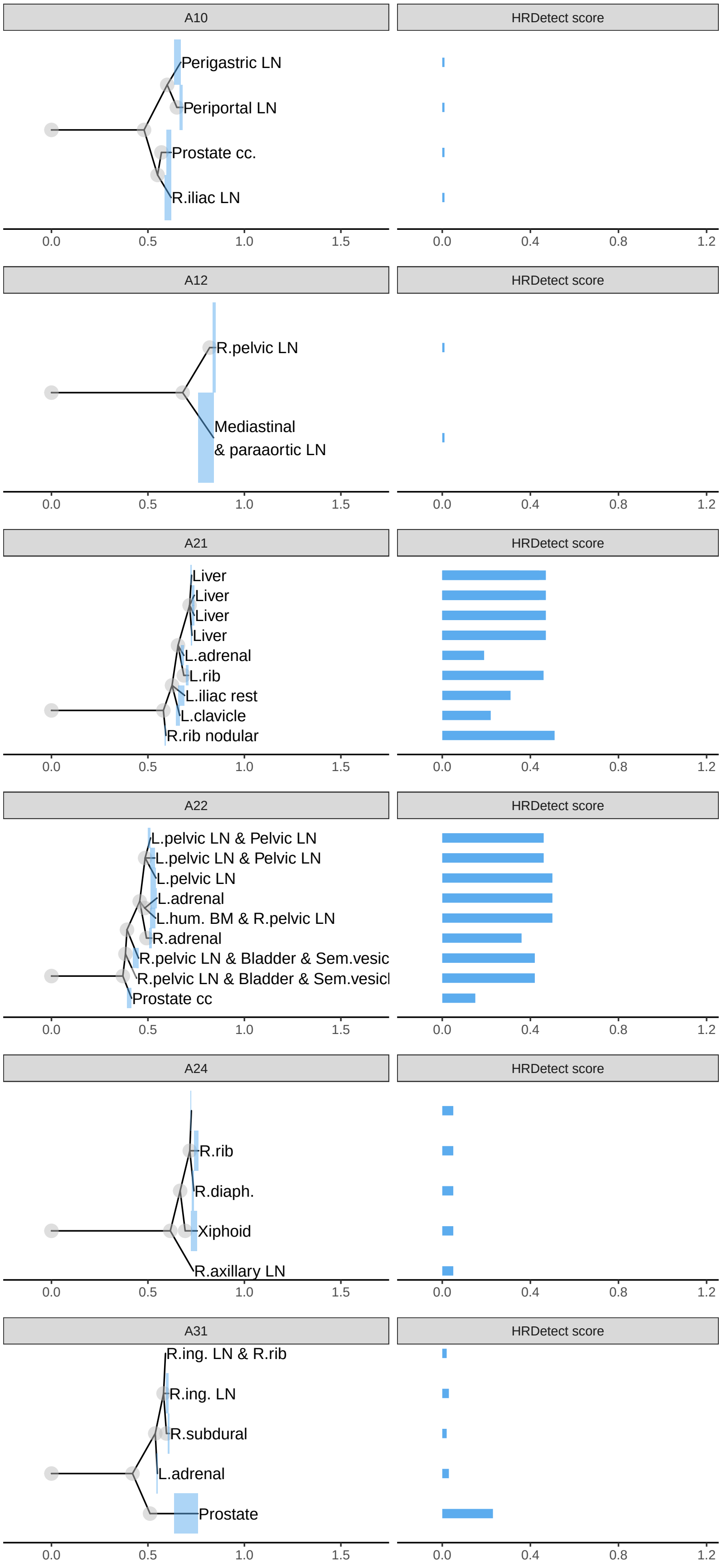
A10
HRDetect score
Perigastric LN
Periportal LN
Prostate cc.
R.iliac LN
0.0 0.5 1.0 1.5
0.0 0.4 0.8 1.2
A12
HRDetect score
R.pelvic LN
Mediastinal
& paraaortic LN
0.0 0.5 1.0 1.5
0.0 0.4 0.8 1.2
A21
HRDetect score
Liver
Liver
Liver
Liver
L.adrenal
L.rib
L.iliac rest
L.clavicle
R.rib nodular
0.0 0.5 1.0 1.5
0.0 0.4 0.8 1.2
A22
HRDetect score
L.pelvic LN & Pelvic LN
L.pelvic LN & Pelvic LN
L.pelvic LN
L.adrenal
L.hum. BM & R.pelvic LN
R.adrenal
R.pelvic LN & Bladder & Sem.vesic
R.pelvic LN & Bladder & Sem.vesicl
Prostate cc
0.0 0.5 1.0 1.5
0.0 0.4 0.8 1.2
A24
HRDetect score
R.rib
R.diaph.
Xiphoid
R.axillary LN
0.0 0.5 1.0 1.5
0.0 0.4 0.8 1.2
A31
HRDetect score
R.ing. LN & R.rib
R.ing. LN
R.subdural
L.adrenal
Prostate
0.0 0.5 1.0 1.5
0.0 0.4 0.8 1.2

**(C)**

### 10 Supplementary Figure 9, Effect of HRDetect score on survival after excluding germline BRCA-mutant cases

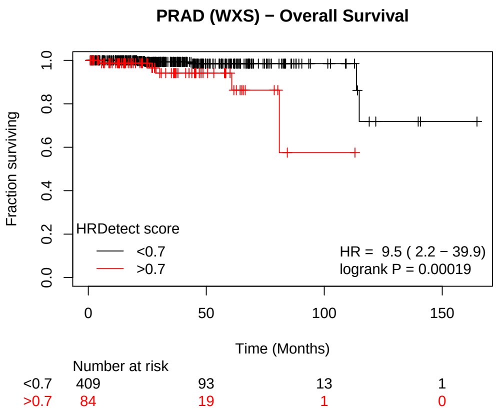


**Suppl.Fig. 9:** HRDetect-score and prognosis, after excluding germline BRCA-mutant cases, TCGA, WES cohort

#### 11 Supplementary Figure 10, Biallelic loss of HR-related genes in the WGS cohorts

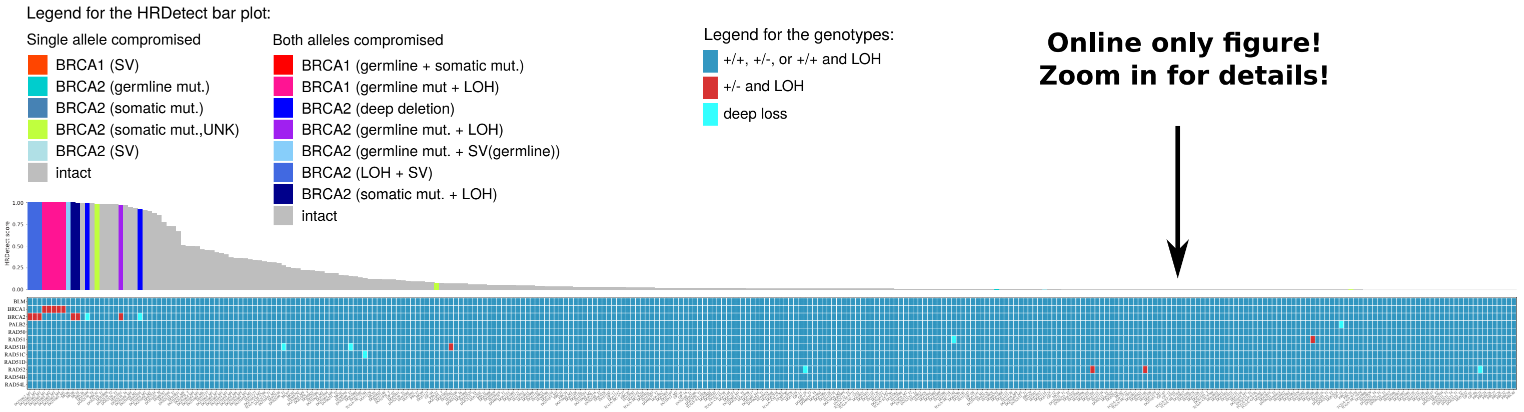


**Suppl.Fig. 10**

#### 12 Supplementary Figure 11, Biallelic loss of HR-related genes in the WES cohort

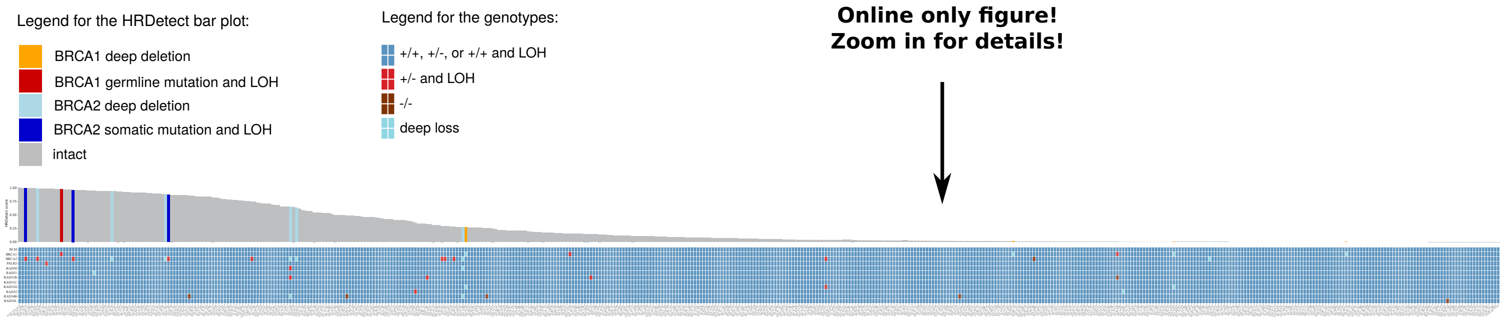


**Suppl.Fig. 11**
